# New Music System Reveals Spectral Contribution to Statistical Learning

**DOI:** 10.1101/2020.04.29.068163

**Authors:** Psyche Loui

## Abstract

Knowledge of speech and music depends upon the ability to perceive relationships between sounds in order to form a stable mental representation of statistical structure. Although evidence exists for the learning of musical scale structure from the statistical properties of sound events, little research has been able to observe how specific acoustic features contribute to statistical learning independent of the effects of long-term exposure. Here, using a new musical system, we show that spectral content is an important cue for acquiring musical scale structure. In two experiments, participants completed probe-tone ratings before and after a half-hour period of exposure to melodies in a novel musical scale with a predefined statistical structure. In Experiment 1, participants were randomly assigned to either a no-exposure control group, or to exposure groups who heard pure tone or complex tone sequences. In Experiment 2, participants were randomly assigned to exposure groups who heard complex tones constructed with odd harmonics or even harmonics. Learning outcome was assessed by correlating pre/post-exposure ratings and the statistical structure of tones within the exposure period. Spectral information significantly affected sensitivity to statistical structure: participants were able to learn after exposure to all tested timbres, but did best at learning with timbres with odd harmonics, which were congruent with scale structure. Results show that spectral amplitude distribution is a useful cue for statistical learning, and suggest that musical scale structure might be acquired through exposure to spectral distribution in sounds.

## 2 Introduction

Implicit learning is a human capacity that is crucial for successful interactions within one’s environment, including via language and music. Abundant evidence has shown that humans can learn about the statistical distribution of sounds from passive exposure to sound sequences (e.g. (Bigand, Perruchet, & Boyer, 1998; Saffran, Aslin, & Newport, 1996; Saffran, Johnson, Aslin, & Newport, 1999). The role of implicitly acquired knowledge for musical structure has been robustly demonstrated in multiple behavioral paradigms (e.g. (Krumhansl, 1990; Loui & Wessel, 2007; Tillmann & McAdams, 2004) and others) and with electrophysiological and neuroimaging indices, even among people who have received no explicit musical training (e.g. (Koelsch, Gunter, Friederici, & Schroger, 2000; Landau & D’Esposito, 2006), among others). This robust evidence for musical knowledge without explicit instruction suggests that music can be a valuable model system that provides a window into how the human mind implicitly acquires knowledge from exposure.

Statistical learning in music is thought to share some mechanisms of learning and memory with language acquisition (McMullen & Saffran, 2004). Computational, behavioral and neurophysiological studies have shown that statistical learning underpins the knowledge-based expectations for musical structures (Pearce, Ruiz, Kapasi, Wiggins, & Bhattacharya, 2009), which bestow musical experiences with emotion and meaning (Meyer, 1956). In assessing implicitly-acquired musical knowledge, a well-replicated behavioral technique is the probe-tone method (Krumhansl & Shepard, 1979), which has been described as a functional hearing test of musical knowledge (Russo, 2009). The probe-tone method involves presenting a musical context (such as a tone sequence) followed by a single tone (i.e. probe tone) to human listeners, who then rate how well the probe tone fits the context. Probe-tone profiles in Western musical scales reflect the frequency of pitch classes in the common-practice Western tonal system, even among untrained listeners (Krumhansl, 1991). This correspondence suggests that some aspect of the implicit knowledge that is reflected in these probe-tone profiles could be acquired from exposure to music in the Western tonal system in the listeners’ environment. To test the contribution of exposure to implicit knowledge using the probe-tone method, Castellano, Bharucha, and Krumhansl (1984) showed that after listening to North Indian rags, Western listeners made probe-tone ratings that were consistent with the distribution of tones they encountered during exposure, suggesting that the tonal hierarchy could be conveyed by the statistical distribution of tones. Additionally, Krumhansl et al. (2000) obtained probe-tone profiles for North Sami yoiks (a vocal musical tradition of Northern Scandinavia) from Western, Finnish, and Sami listeners, and showed that Finnish listeners’ probe-tone ratings reflected some familiarity of both Western and Yoik scale structures, again suggesting that probe-tone profiles reflect sensitivity to statistical structures of musical sounds in one’s environment.

While these cross-cultural methods provide powerful evidence that knowledge of scale structure is not limited to Western tonal systems, these results could not disentangle long-term musical knowledge (i.e. knowledge that is acquired from birth and accumulated over the lifespan) and short-term statistical learning that might be operating on a moment-by-moment basis. More generally, the question of how rapidly humans could acquire new knowledge about scale structure is difficult to address using conventional musical systems of any culture. This is because conventional musical systems evolved through complex cultural evolution over time (Cross, 2001; Mithen, 2007); thus they are already overlearned throughout the lifespan as a result of exposure within that culture (to music as well as other sounds such as speech) (Patel & Daniele, 2003). Thus, conventional musical systems cannot capture learning de novo in a way that is not intertwined with culture and cultural evolution.

To circumvent this challenge, several studies have turned to test learning of novel musical systems (e.g. (Creel & Newport, 2002); Leung and Dean (2018)). Loui et al (Loui, 2012; Loui & Wessel, 2008; Loui, Wessel, & Hudson Kam, 2010; Loui, Wu, Wessel, & Knight, 2009) developed a musical system that uses the Bohlen-Pierce (BP) scale, which differs from existing musical scales in important ways. While other musical scales are based on the octave, which is a doubling (2:1 ratio) in frequency, the BP scale is based on the 3:1 frequency ratio (tritave). The equal-tempered Western chromatic scale is based on 12 logarithmically-even divisions of the octave; this enables the selection of certain tones, such as steps 0 (starting point), 4, and 7 along the chromatic scale, that approximate the 3:4:5 integer ratio in frequency, low-integer ratios that lead to consonant sounds that form stable chords in traditional tonal harmony. In contrast, the BP scale divides the tritave into 13 logarithmically-even steps, resulting in steps 0, 6, and 10 being stable tones that approximate a 3:5:7 integer ratio, thus forming a chord in the BP scale (Krumhansl, 1987; Mathews, Pierce, Reeves, & Roberts, 1988). Learning of the statistical structure of the music, which was operationalized as the increase in sensitivity to event-probabilities of different pitches, was captured by comparing probe-tone ratings before and after exposure to tone sequences in the BP scale (Loui et al, 2010). Statistical sensitivity was assessed by the correlation between probe-tone ratings and the event distribution of tones in the exposure set, and results showed an increase in correlation from pre-exposure to post-exposure probe-tone ratings, confirming learning of the BP scale as indicated by increased sensitivity to its statistical structure.

Importantly, the tone sequences in the previous studies were pure tones ranging in fundamental frequency from 220 Hz to 660 Hz, with a single acoustic frequency presented for each acoustic event in time. Thus, there was a one-to-one correspondence between spectral information (i.e. acoustic frequency content) and pitch class information. Although pure tones in this range provide a clear percept of pitch, they are not representative of real-world acoustic input because they lack additional acoustic energy along the frequency spectrum (Sethares, 2004). This additional spectral energy provides crucial cues to the listener on many aspects of sound object recognition (Bregman, 1990), including the identity of a musical instrument based on its timbre (Wessel, 1979), the identity of a human speaker based on their voice (Belin, Fecteau, & Bédard, 2004), and the identity of phonemes in speech (Smith, 1951). Spectral information is important for identifying tone sequences even among nonhuman animals such as starlings, who rely on the shape of energy distributed along the frequency spectrum (rather than pitch information per se) to recognize sound patterns (Bregman, Patel, & Gentner, 2016). For periodic sounds, the shape of the spectral distribution is dependent on multiple factors including the frequency of the weighted average of the harmonics (i.e. the spectral centroid), and the spacing between individual frequency components (i.e. spectral fine structure), all of which contribute to the percepts of timbre (Caclin, McAdams, Smith, & Winsberg, 2005; Sethares, 2004; Wessel, 1979).

Despite the importance of spectral information, little is known about the role of spectral information in the statistical learning of scale structure. How does spectral information influence the learning of musical scales? One possibility is that spectral information is orthogonal and unrelated to the event structure of sound sequences. In that case, probe-tone ratings should not differ between different timbres. An alternative hypothesis is that spectral information, as determined here by the spacing between individual harmonics, can be a cue towards the statistical structure of sounds. This latter hypothesis would predict that probe-tone ratings would be better correlated with the scale structure, indicating better learning, if spectral information in its timbre is consistent with the scale structure. The BP scale offers an optimal test for the relationship between spectral information and statistical learning, since most listeners have no experience with this new musical scale.

The hypothesis that spectral information provides cues for the statistical structure of sound sequences makes a strong prediction: that tone sequences in a timbre that is congruent with the musical scale should help in learning the musical scale. Thus, as the BP scale is based on the 3:1 frequency ratio, timbres with harmonics that are spaced apart in 3:1 frequency ratios are congruent with the scale structure, whereas timbres with harmonics that are spaced apart in 2:1 frequency ratios are incongruent with the BP scale structure (despite being congruent with the Western scale structure). By generating complex tones that vary in specific placement of their harmonically related partials (Shepard, 1964), it is possible to manipulate the spacing between harmonics to be congruent or incongruent with the BP scale (see Figure 1), thus directly testing the effect of spectral information on statistical learning.

**Figure 1.**
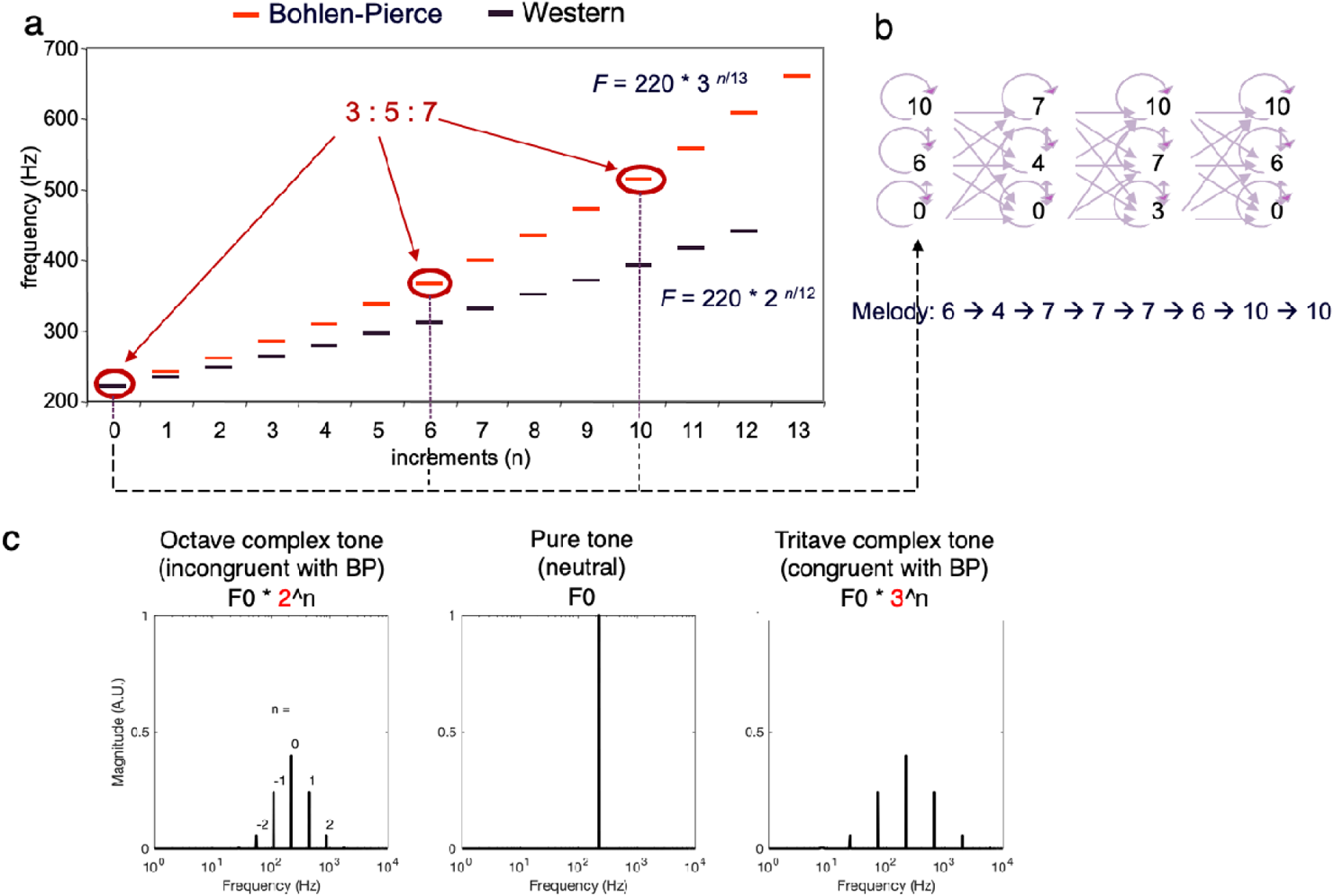
Properties of the Bohlen-Pierce scale motivate stimuli generation in Experiment 1. **a)** The BP scale is based on a 3:1 ratio, unlike Western scale which is based on 2:1 ratio. Tones along the BP scale that approximate low-integer ratios in frequency (3:5:7) were chosen as chords in the new scale. **b)** Four chords, each containing tones that thus approximated low-integer ratios, were strung together to form a chord progression, and used to generate tone sequences according to a finite-state grammar. Purple arrows show the paths of possible movement between tones in the finite-state grammar, along with an example of a resultant melody, which is a legal string in this finite-state grammar. **c)** Frequency-domain representations of sound stimuli used in Experiment 1: octave-based complex tones (partials spaced apart in 2:1 frequency ratios), pure tones (no partials), and tritave-based complex tones (partials spaced apart in 3:1 frequency ratios).

Here we test the role of spectral content on statistical learning, by comparing learning of the BP scale between timbres that are spectrally congruent and incongruent with the scale. Learning is quantified by improvements in the accuracy of probe-tone ratings as a result of exposure to tone sequences in the BP scale. In Experiment 1, we compared learning among participants who heard pure tones (no harmonic partials), tritave-based complex tones (complex tones where partials were related to the fundamental in 3:1 multiples in frequency), and octave-based complex tones (complex tones with partials related to the fundamental in 2:1 multiples of frequency), and a no-exposure control. Since the tritave complex tones are congruent with the scale structure, whereas the octave complex tones are incongruent with the scale structure, we predict that learning would be best with tritave complex tones, and worse with octave complex tones.

In Experiment 2, we compared learning among participants who heard odd harmonics and even harmonics, with the prediction that odd harmonics would help learning better as they were more spectrally congruent with the BP scale due to its 3:5:7 (odd numbers) integer frequency ratio. Together, these experiments test the role of spectral content in the acquisition of musical knowledge, which informs the ability to learn from sound input more generally.

## 3 Experiment 1

### 3.1 Materials and Methods

#### 3.1.1 Participants

Ninety-six undergraduates (57 females, 37 males) at the University of California at Berkeley participated in this experiment in return for course credit. Participants were on average 20.79 years of age (SD = 4.32 years), and all participants reported having normal hearing. The sample size was chosen to be the same as previous experiments that had used the same experiment design to assess statistical learning of the BP scale (Loui & Schlaug, 2012; Loui & Wessel, 2008; Loui, Wessel, & Hudson Kam, 2006; Loui et al., 2010). Since our previous work had shown that participants could learn the BP scale regardless of their amount of previous musical training (Loui et al., 2010), we enrolled participants regardless of how much musical training they had received. Of the resulting sample, n = 23 reported having no musical training. The rest reported an average of 6.43 years of musical training (SD = 5.18 years) in a variety of instruments including piano (n = 40), violin (n = 14), guitar (n = 12), clarinet (n = 6), flute (n = 6), cello (n = 4). Each participant was randomly assigned to an exposure condition (n = 24 participants per exposure condition). After providing written informed consent, participants were tested in a sound-attenuated room while facing a Dell desktop PC running Max/MSP software (Zicarelli, 1998). Stimuli were presented at approximately 70 dB through Sennheiser HD 280 headphones. All data and code used in analysis are available on https://osf.io/pjkq2/.

#### 3.1.2 Procedure

The experiment was conducted in three phases: 1) pre-exposure probe-tone ratings test, 2) exposure phase, and 3) post-exposure probe-tone ratings test.

##### Pre-exposure probe-tone ratings test

Thirteen trials were conducted in this phase. In each trial, participants were presented with a melody in the Bohlen-Pierce scale, followed by a tone (Krumhansl, 1991). The melody was 4 seconds long, consisting of 8 tones of 500 ms each, including rise and fall times of 5 ms each. The probe tone began 500 ms after the end of the melody, and was also 500 ms including rise and fall times of 5 ms each. Participants’ task was to rate how well the last tone (i.e. the probe tone) fit the preceding melody, on a scale of 1 (least fitting) to 7 (best fitting).

##### Exposure phase

Participants were presented with 400 melodies in the BP scale. Each melody was 4 seconds long, consisting of 8 tones of 500 milliseconds each, including rise and fall times of 5 ms each, with a 500 ms silent gap between successive melodies, resulting in an exposure phase that lasted 30 minutes. The pitches of the melodies were determined by a finite-state grammar previously described in Loui et al. (2010) and illustrated in Figure 1a-b: Tones along the BP scale that approximated low-integer ratios in frequency (3:5:7 as mentioned in Introduction, see Figure 1a) were chosen as chords in the new scale. Four chords, each containing tones that thus approximated low-integer ratios, were strung together to form a chord progression. Applied to finite-state grammar terms, each chord represented a state of the finite-state grammar. Each tone in each chord could either repeat itself, or move within other tones in the same chord, or move forward to the next chord. Figure 1b shows the paths of possible movement between tones in this finite-state grammar, along with an example of a resultant melody, which is a legal string in this finite-state grammar. The melodies were presented in one of four possible timbre conditions as described below: congruent (Tritave complex tones), incongruent (Octave complex tones), or neutral (Pure tones), or the no-exposure control condition. Figure 1c shows a schematic of how the Tritave complex tones, the Octave complex tones, and the pure tones were generated.

###### Neutral Condition

**Pure tones** were computer-generated with fundamental frequency only, i.e. no partials. In this condition, the only frequency was the fundamental frequency (F0) as specified by the BP scale. Thus these tones provided a clear percept of pitch, but did not provide any spectral cues as to the tuning system.

###### Congruent Condition

**Tritave complex tones** were computer-generated complex tones with five partials centering around the spectral centroid, where the partials were related to the fundamental in 3:1 ratios (tritave) in frequency. The spectral centroid was the same as the pure tone F0. The amplitudes of frequency components of the complex tone were scaled with a normal curve centering around the spectral centroid. Thus, the five frequency components were .11 (3^-2), .33 (3^-1), 1 (3^0, same as the spectral centroid), 3 (3^1), and 9 (3^2) times the F0. Since the BP scale is based on the 3:1 frequency ratio (instead of 2:1 frequency ratio, i.e. the octave), the timbre of tones in this condition is congruent with a tritave-based musical system.

###### Incongruent Condition

**Octave complex tones** were computer-generated complex tones with five partials centering around the target frequency, where the partials were related to the fundamental in 2:1 ratios in frequency. The spectral centroid was again the same as the pure tone F0. As above, the amplitudes of frequency components of the complex tone were scaled with a normal curve centering around the spectral centroid. However, here the five frequency components were .25 (2^-2), .5 (2^-1), 1 (2^0, same as the spectral centroid), 2 (2^1), and 4 (2^2) times the F0. Thus these tones were incongruous with the BP scale, but consistent with Western and other musical scales that are based on the 2:1 frequency ratio.

###### Control Condition

In a **no-exposure control condition**, participants made probe-tone ratings twice, using the same procedures as the Neutral (pure tone) condition. They were not given exposure to any auditory stimuli between the pre- and post-exposure ratings; instead, they were asked to sit quietly for 30 minutes between the two probe-tone tests, as this was the duration of the exposure condition.

##### Post-exposure probe-tone ratings test

Probe-tone ratings were conducted again after exposure, using the same methods as phase 1.

### 3.2 Results

Figure 2 shows the probe-tone ratings before and after exposure for each condition. For all conditions except the no-exposure control, the red line (post-exposure ratings) is visibly more correlated with the blue line (exposure distribution) compared to the black line (pre-exposure ratings), suggesting an increase in correlations overall and consistent with acquisition of the statistical structure of the BP scale following exposure.

**Figure 2.**
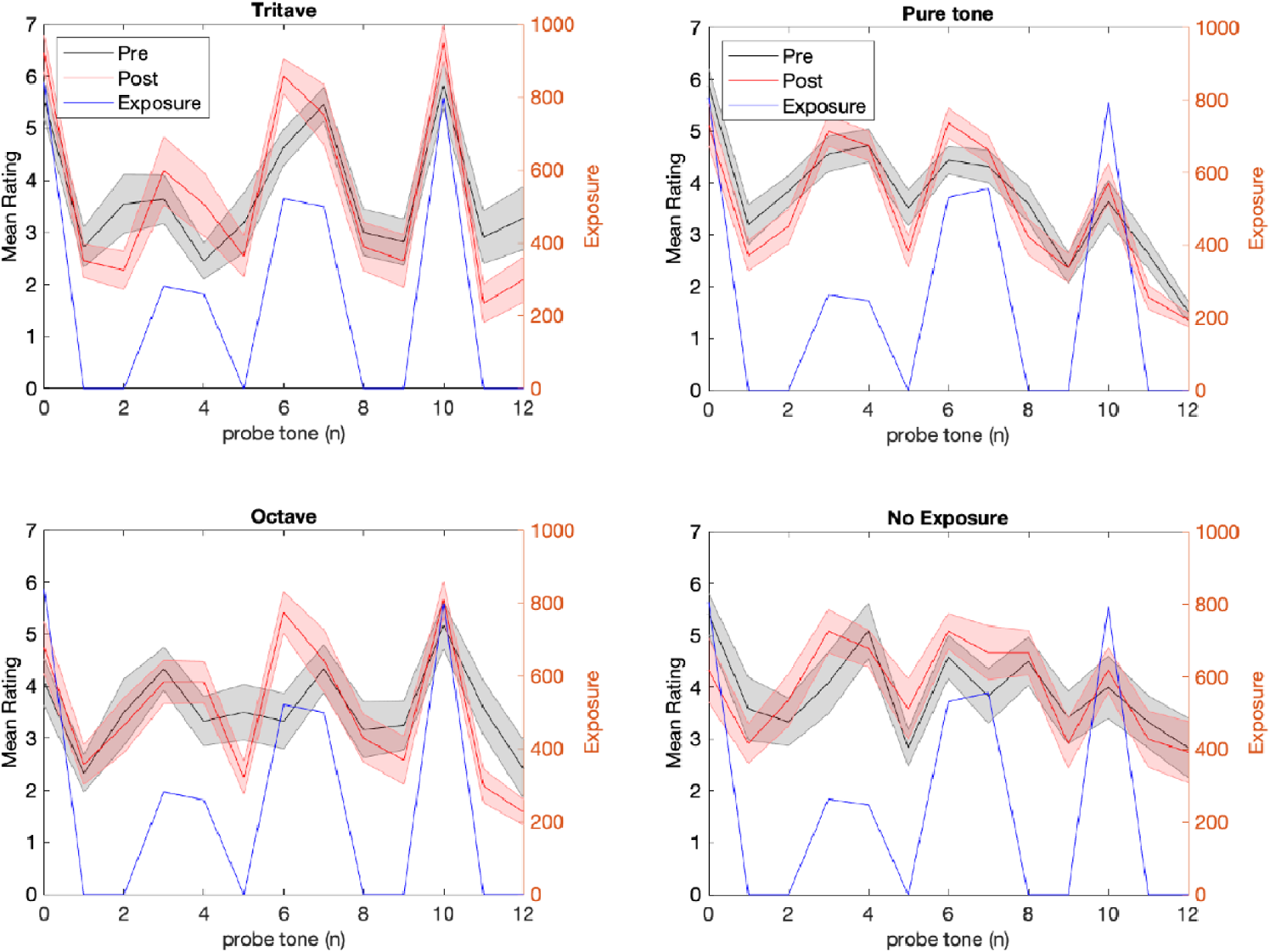
Probe-tone ratings in the four experimental conditions. X-axis represents the probe tone in steps along the BP scale. Pre-exposure ratings are in black and post-exposure ratings are in red. The exposure profile is shown in blue. The shaded error bars represent ±1 between-subject standard error. The red ratings are more highly correlated with the blue exposure profile than the black ratings (except for the no-exposure condition), suggesting that learning occurred as a result of exposure.

Sensitivity to statistical structure of the scale was quantified by the correlation between probe-tone ratings and exposure. Figure 3a shows the Pearson correlation between exposure profile and probe tone ratings, computed for each subject, for all four conditions. These correlations are higher post-exposure than pre-exposure in all exposure conditions, but not in the no-exposure control condition.

**Figure 3.**
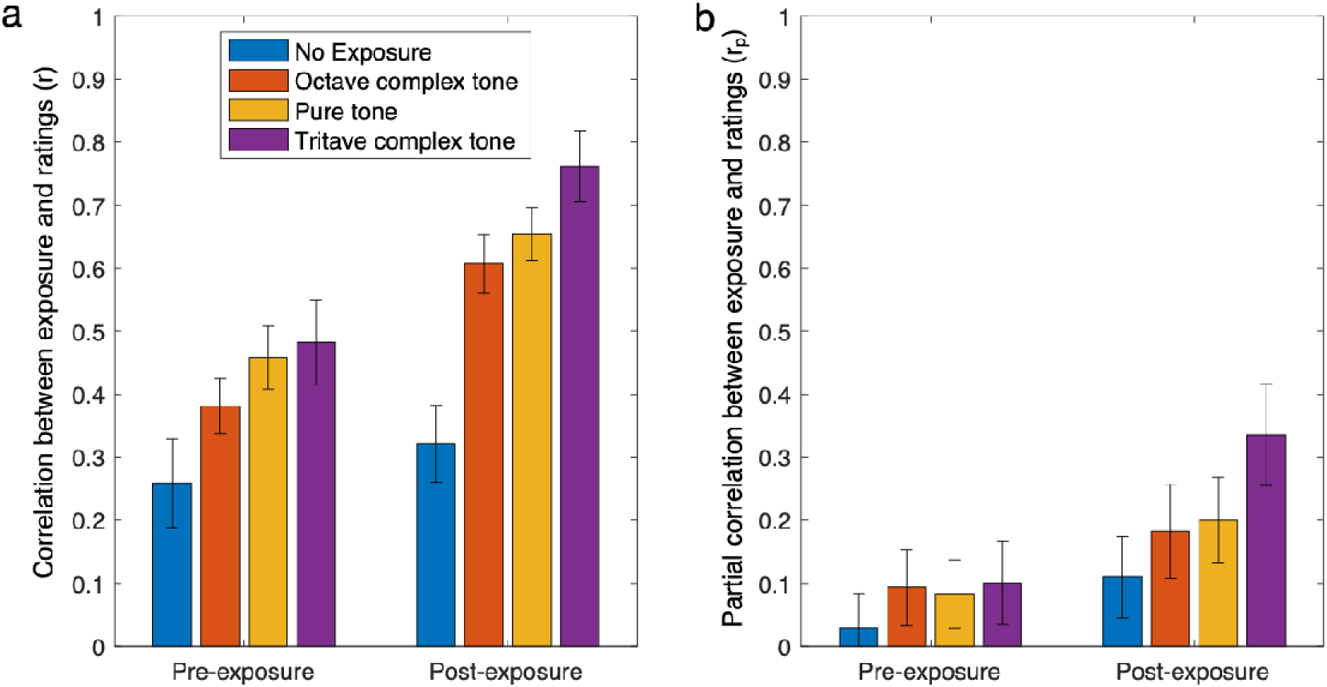
Correlations **(a)** and partial correlations **(b)** between probe-tone ratings and exposure frequencies. Error bars show between-subject standard error.

As shown by Figure 3a, pre-exposure correlations were above zero. This was likely because participants were influenced by the short-term exposure of the melody that was used to obtain the probe-tone ratings. Since a probe-tone context had to be presented in order to obtain ratings, participants relied on the context to make ratings before exposure. To disentangle the effect of the context from the effect of exposure on probe-tone ratings, partial correlations were obtained by partialling out the event distribution of the melody from the relationship between the ratings and the event distribution of the exposure. Figure 3b shows the partial correlations for each condition. Pre-exposure ratings were close to zero whereas post-exposure ratings were higher and above zero, except for the no-exposure control condition. As the correlations and partial correlations were not normally distributed (Kolmogorov-Smirnov tests p < .05), they were Fisher-transformed to z-values to transform the data towards a normal distribution prior to statistical testing. A mixed-effects multivariate ANOVA was conducted with the dependent variables of the z-transformed correlations and partial correlations, and a within-subjects factor of exposure (pre-exposure vs. post-exposure) and a between-subjects factor of condition (Tritave, Octave, Pure tone, No exposure). Table 1 shows the results of this multivariate ANOVA, showing a significant between-subjects effect of condition (F(6,182) = 4.38, p < .001, partial η^2^ = .13), confirming that participants’ ratings differed between groups. There was also a significant within-subject effect of exposure (F(2,90) = 32.50, p < .001, partial η^2^ = .42), confirming that exposure affected ratings. Importantly, there was a significant interaction between exposure and condition (F(6,192) = 2.94, p = .009, partial η^2^ = .088), confirming that exposure affected the different groups differently.

**Table 1.** Multivariate ANOVA results from Experiment 1. **a)** Mixed effects MANOVA with all four groups included. **b)** MANOVA results from Experiment 1, separately showing results for the no-exposure conditions and the three exposure conditions.

| a) Effect | | F | Hypothesis df | Error df | p | Partial $\eta^2$ |
| --- | --- | --- | --- | --- | --- | --- |
| Between Subjects | Intercept | 177 | 2 | 90 | .001 | 0.80 |
|  | Condition | 4.38 | 6 | 182 | .001 | 0.13 |
| Within Subjects | Exposure | 32.50 | 2 | 90 | .001 | 0.42 |
|  | Exposure * Condition | 2.94 | 6 | 182 | .009 | 0.088 |

| b) | Effect | | F | Hypothesis df | Error df | p | Partial $\eta^2$ |
| --- | --- | --- | --- | --- | --- | --- | --- |
| No exposure | Between Subjects | Intercept | 13.14 | 2 | 22 | <.001 | 0.54 |
|  | Within Subjects | Exposure | 1.25 | 2 | 22 | 0.31 | 0.10 |
| Exposure | Between Subjects | Intercept | 190.42 | 2 | 67 | <.001 | 0.85 |
|  |  | Condition | 2.02 | 4 | 136 | 0.096 | 0.056 |
|  | Within Subjects | Exposure | 34.562 | 2 | 67 | <.001 | 0.51 |
|  |  | Exposure * Condition | 1.56 | 4 | 136 | 0.188 | 0.044 |

While the above analysis suggested that our different exposure conditions affected learning, the no-exposure group was included in the above analysis, and could have driven the interaction, as participants who received the no-exposure condition were not expected to learn from exposure. To separate the no-exposure group from the other groups who received exposure, follow-up multivariate ANOVAs were conducted separately for the no-exposure group and the other exposure groups, with the dependent variables of the z-transformed correlations and partial correlations, and a within-subjects factor of exposure (pre-exposure vs. post-exposure), and an additional between-subjects factor of condition (Tritave, Octave, Pure tone). The no-exposure group did not show an effect of exposure (F(2,22) = 1.25, p - .31, partial η^2^, = .1), as expected.

The three groups that received Tritave, Octave, and Pure tone exposures showed significant effects of exposure (F(2,67) = 34.56, p < .001, partial η^2^, = .51), and a marginally significant effect of condition (F(4,136) = 2.02, p = .096, partial η^2^, = .056). The interaction between exposure and condition was no longer significant once the no-exposure condition was removed (F(4,136) = 1.56, p = 0.188, partial η^2^, = .044), suggesting that the previously-observed effect of condition and interaction between exposure and condition was indeed driven by the no-exposure condition being different from the other groups.

To follow up on the effects of exposure of each exposure condition, and to treat the no-exposure condition as a true control condition, a Tukey’s HSD test was applied to compare the z-transformed correlations for all the exposure conditions against the no-exposure control, while correcting for the family-wise error rate. Table 2a shows the results of this pairwise comparison. Results show that all groups learned significantly better than the no-exposure group as measured by the z-transformed correlation scores. This confirms, as expected, that all groups who received exposure showed significantly more accurate ratings than the group that received no exposure.

**Table 2.**
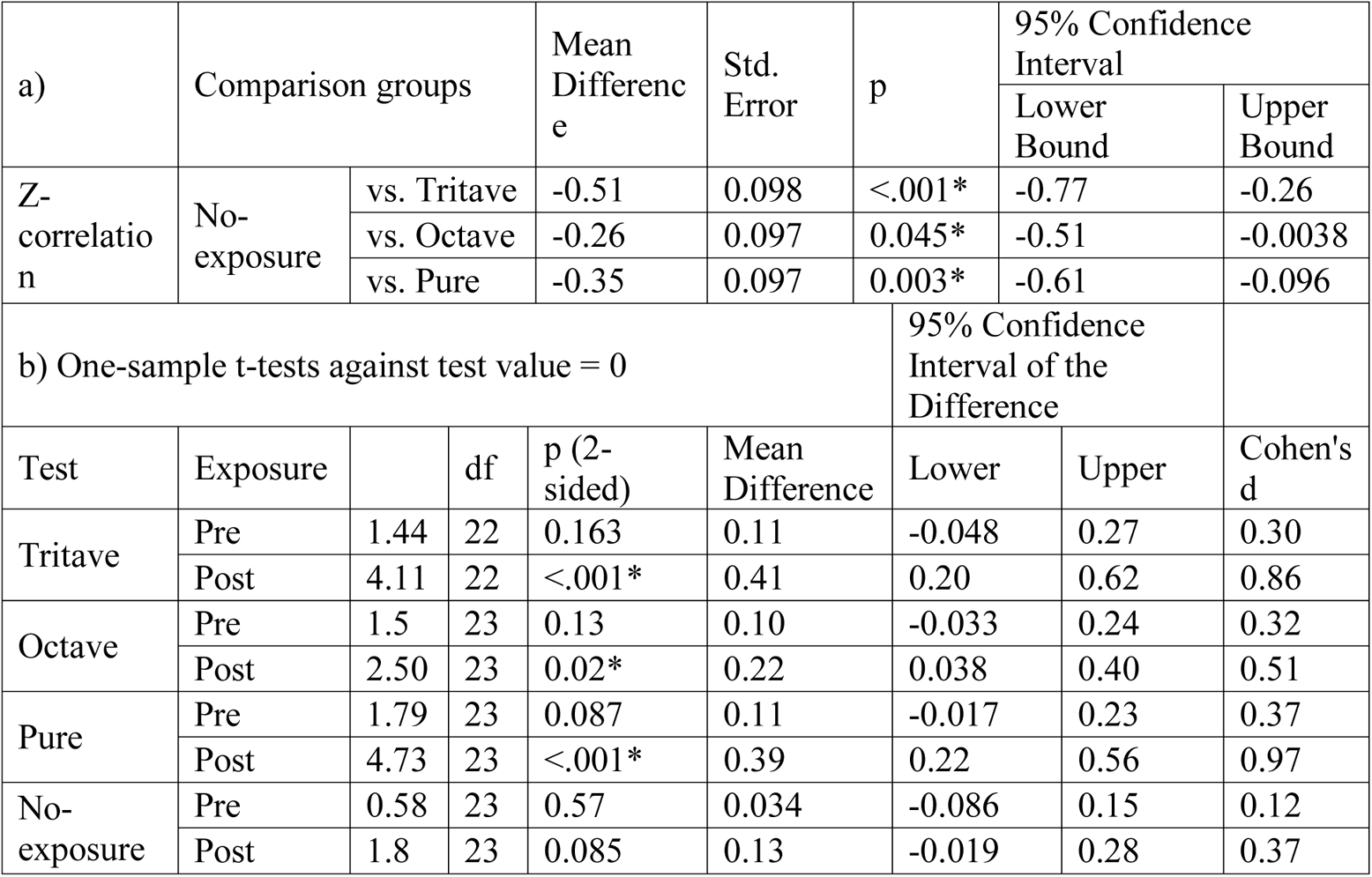
Pairwise comparisons between groups. **a)** Pairwise comparisons of z-transformed correlations for each exposure condition against no-exposure control. * denotes significant difference from no-exposure control group at p < .05, Tukey’s HSD test. **b)** Two-tailed t-tests of z-transformed pre- and post-exposure partial correlations against the chance level of zero. * denotes significant deviation at p < .05 from chance level of 0, two-tailed t-test.

While the correlations were used to compare against a no-exposure control, the partial correlations were tested against a chance level of zero (no correlation between ratings and exposure), which would indicate random responding. Table 2b shows two-tailed t-tests of z-transformed partial correlations against chance level. The Tritave, Octave, and Pure tone post-exposure conditions, i.e. all the post-exposure conditions, but not the pre-exposure or the no-exposure control condition, showed significantly above-chance partial correlations. This confirmed that participants acquired the scale structure as a result of exposure, resulting in non-random ratings after exposure that were not explained by the melody used in the ratings task.

### 3.3 Discussion

Experiment 1 showed statistical learning of the BP scale structure in all exposure conditions, and not in the no-exposure condition, as expected. Although the effect of condition was significant, as was the effect of exposure and the interaction between exposure and condition, this was driven by the no-exposure condition showing different results from the other groups. Treating the no-exposure condition as a true control, pairwise comparisons showed that each exposure condition resulted in better performance than the no-exposure control. Each exposure condition also resulted in significantly above-chance partial correlations after exposure. These results suggest robust learning from exposure. Although the Tritave complex tone condition resulted in higher correlations and higher partial correlations than the pure tone and Octave complex tone conditions, as seen in Figure 2, the effect of spectral distribution, as manipulated by these complex tones, was only marginally significant. This could be because the experiment was underpowered due to the relatively small sample size: the sample size of this experiment was chosen based on previous studies which had shown statistical learning (as stated in Methods), but previous studies had not investigated the effect of spectral information on learning. Another possibility was that the manipulation of spectral content was not strong enough. As the Tritave and Octave complex tones contained only five frequency components each, this resulted in a relatively sparse spectrum with only fairly limited spectral information. Furthermore, the frequency components had to be spaced further apart in Tritave complex tones than in Octave complex tones, due simply to the fact that a 3:1 ratio is always larger than a 2:1 ratio. It could be the case that the spectrum was too sparse to be informative, i.e., more spectral information was necessary to provide a stronger test of the effect of spectral information on statistical learning.

## 4 Experiment 2

Experiment 1 showed conclusive evidence for learning from exposure, replicating previous studies (Loui & Schlaug, 2012; Loui & Wessel, 2008; Loui et al., 2006; Loui et al., 2010; Loui et al., 2009), but only weak evidence for better performance for the Tritave than for the Octave complex tone in learning the BP scale. Results from Experiment 1 could also have been confounded by concerns of sample size, and of participants being more familiar with octave spacing among frequency components in natural sounds. To rule out these possibilities, for Experiment 2 we generated novel tones with odd and even harmonics. Chords in the BP scale structure are determined by odd-number low-integer ratios 3:5:7 (see Figure 1a). We reasoned that if the congruence between spectrum and scale structure enabled better statistical learning, then tones with frequencies that obey 3:5:7 integer ratios, i.e. tones that have energy at odd integer harmonics in their spectrum, would be predicted to enable better learning, compared for example to even-integer harmonics (see Figure 4a-b). Some support for this comes from the observation that in the small existing repertoire of art music written in the BP scale, the majority is written for instruments that play in odd harmonics due to their closed-tube physical structures, such as the clarinet, the bass clarinet, and the pan flute (Advocat, 2010; Hajdu & Didkovsky, 2018). Here we compare statistical learning of the same BP scale system in complex tones with odd-integer harmonics versus even-integer harmonics. Odd harmonics are predicted to be more conducive to learning.

**Figure 4.**
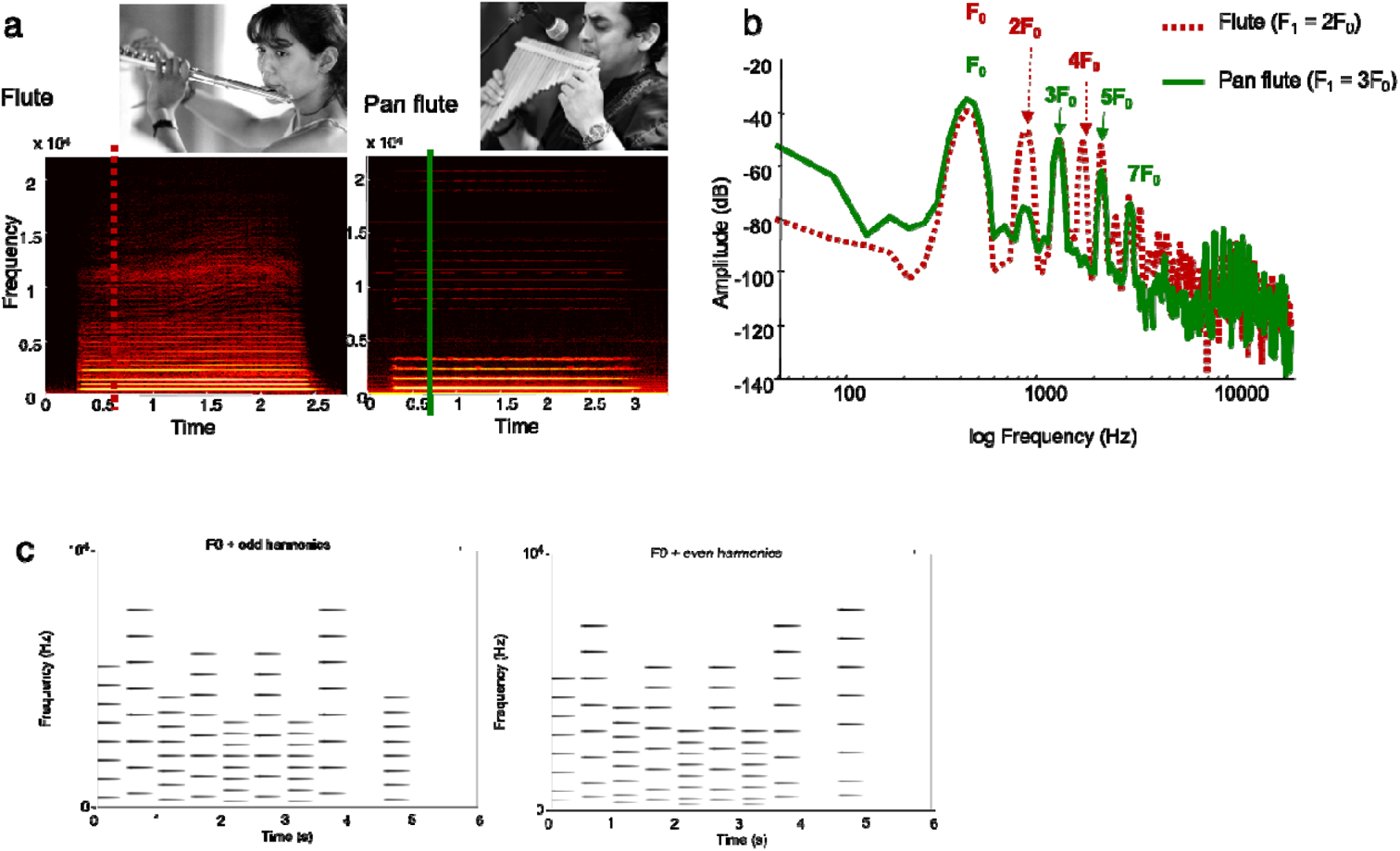
Comparing timbres with odd and even harmonics. **a)** Physical shape and representative spectrograms of an A4 played on the flute and the pan flute. Sounds of the pan flute, due to its closed tube structure, have more energy at odd-integer harmonics, compared to the sounds of the Western flute, which has an open-tube structure and thus has energy at both odd and even harmonics. **b)** A comparison of the spectral distribution of a single note (A4) played by a flute (red dotted line) and a pan flute (green solid line). While both instruments show energy at the F0 of 440 Hz, the flute shows overtones at 2, 3, 4, and 5 x the F0. In contrast, the pan flute shows energy at 3, 5, and 7 x the F0. In other words, the pan flute emphasizes odd integer multiples of the F0 whereas the Western flute emphasizes both odd and even integer multiples. c) Spectrogram of one representative trial of the current Experiment 2. Both spectrograms show a probe tone trial, where a single melody in the BP scale is presented, followed by a pause, and a final tone. Left panel shows the odd harmonics condition, where each F0 is accompanied by harmonics at 3, 5, 7, 9, 11, 13, and 15 times the F0. Right panel shows the even harmonics condition, where each F0 is accompanied by harmonics at 2, 4, 6, 8, 10, 12, and 14 times the F0. The odd harmonics condition is predicted to lead to better learning.

### 4.1 Materials and Methods

#### 4.1.1 Participants

126 young adults (56 females, 70 males) participated in this experiment in return for payment or course credit. A power analysis was conducted using effect sizes from Experiment 1 on the effect of condition (partial η^2^ of 0.056) and the interaction between exposure and condition (partial η^2^ of 0.044), to detect a significant effect of condition and a significant exposure-condition interaction at an alpha level of 0.05 in two groups. The power calculation yielded a necessary n of 40 per condition for a significant effect of condition with 85% power, and an n of 52 per condition for a significant interaction between exposure and condition with 85% power. Participants were on average 24.1 years of age (SD = 3.18 years), and all participants reported having normal hearing. N = 61 reported having no musical training. The rest reported an average of 6.43 years of musical training (SD = 5.18 years) in a variety of instruments including piano (n = 46), guitar (n = 13), violin (n = 12), flute (n = 7), saxophone (4), and other instruments (cello, percussion, voice, n = 1 each). Each participant was randomly assigned to an odd or even harmonics exposure condition, resulting in n = 63 for each group. The study began in-person in a testing room on campus at Northeastern University; however due to the Covid-19 pandemic the study moved online after an online testing protocol was approved by the IRB of Northeastern University. This resulted in N = 26 (13 in the odd condition and 13 in the even condition) who were tested in-person, and N = 100 (50 in the odd condition and 50 in the even condition) who were tested online using Amazon’s Mechanical Turk. To mitigate concerns over sound quality for online experiments, a headphone screening was added to the online study (Woods, Siegel, Traer, & McDermott, 2017), and only participants who scored 5/6 or above and had no missing data were analyzed to ensure data quality. Results were then aggregated between in-person and online studies. Examples of stimuli, all data, and code used in analysis are available on https://osf.io/pjkq2/.

#### 4.1.2 Procedure

The experiment was conducted in three phases, same as Experiment 1: 1) pre-exposure probe-tone ratings test, 2) exposure phase, and 3) post-exposure probe-tone ratings test. However, in Experiment 2 there were only two groups: Odd harmonics and Even harmonics. Participants were randomly assigned into one of the two groups, thus being exposed to the BP scale in either even-integer harmonics or odd-integer harmonics.

##### Pre-exposure probe-tone ratings test

Thirteen trials were conducted in this phase. In each trial, participants were presented with a melody in the Bohlen-Pierce scale, followed by a tone (Krumhansl, 1991). The melody was 4 seconds long, consisting of 8 tones of 500 ms in inter-onset intervals. Each tone was 425 ms in overall duration, including a rise time of 75 ms and a fall time of 350 ms. This amplitude envelope was slightly different from Experiment 1 to sound more pleasant compared to tones with a flat amplitude envelope from Experiment 1. The probe tone began 500 ms after the end of the melody, and was also 425 ms including rise time of 75 ms and fall time of 350 ms. Participants’ task was to rate how well the last tone (i.e. the probe tone) fit the preceding melody, on a scale of 1 (least fitting) to 7 (best fitting).

##### Exposure phase

Participants were presented with 400 melodies in the BP scale. The F0s of the melodies were from the same finite-state grammar system as described in Experiment 1. Each melody was 4 seconds long, consisting of 8 tones of 500 ms in inter-onset time, with each tone lasting 425 ms including rise times of 75 ms and fall times of 350 ms. A 500 ms silent gap was inserted between successive melodies, resulting in an exposure phase that lasted 30 minutes. The melodies were presented in one of two possible timbre conditions as described below: Odd Harmonics and Even Harmonics. Figure 4c shows spectrograms of a single representative trial of the probe tone ratings test, in the odd harmonics condition (left) and in the even harmonics condition (right). Task instructions explained that participants may relax or do their own work during this time, but may not use their headphones, or talk, or listen to other sounds during the half-hour exposure.

##### Odd Harmonics

were computer-generated complex tones with an F0 in the BP scale as determined by the finite-state grammar, and 7 additional overtones that were 3, 5, 7, 9, 11, 13, and 15 times the F0. All the overtones were the same in amplitude as the F0.

##### Even Harmonics

were computer-generated complex tones with an F0 in the BP scale as determined by the finite-state grammar, and 7 additional overtones that were 2, 4, 6, 8, 10, 12, and 14 times the F0. All the overtones were the same in amplitude as the F0.

##### Post-exposure ratings test

were conducted again after exposure, using the same methods as the pre-exposure ratings described above.

### 4.2 Results

Figure 5a shows the probe-tone ratings before and after exposure, for the odd and even harmonics conditions. For both the odd and even harmonics conditions, the red line (post-exposure ratings) was visibly more correlated with the blue line (exposure distribution) compared to the black line (pre-exposure ratings), suggesting an increase in correlations overall and consistent with acquisition of the statistical structure of the BP scale following exposure. This correlation is computed for each subject and shown in Figure 5b, which shows the Pearson correlation between exposure profile and probe tone ratings for the odd and even harmonics conditions (left), and the partial correlations from the same data (right) after the event frequencies of the melody used to obtain the probe tone ratings were partialled out. Ratings from the odd harmonics condition showed higher correlation with exposure than the even harmonics condition, suggesting that participants were more sensitive to the scale when listening to odd harmonics both before and after exposure. When the effects of the melody used to obtain the probe tone ratings were partialled out, the pre-exposure partial correlations were close to zero for both odd and even harmonics, suggesting that some of the sensitivity as shown by the bivariate correlations was explained by the melody used to obtain the ratings. Importantly for the partial correlations, only the odd harmonics condition showed an improvement after exposure and above-zero correlation post-exposure, whereas the even harmonics condition did not show an improvement in correlation after exposure, suggesting that participants who heard the even harmonics were not able to learn from exposure as well as those who heard the odd harmonics.

**Figure 5.**
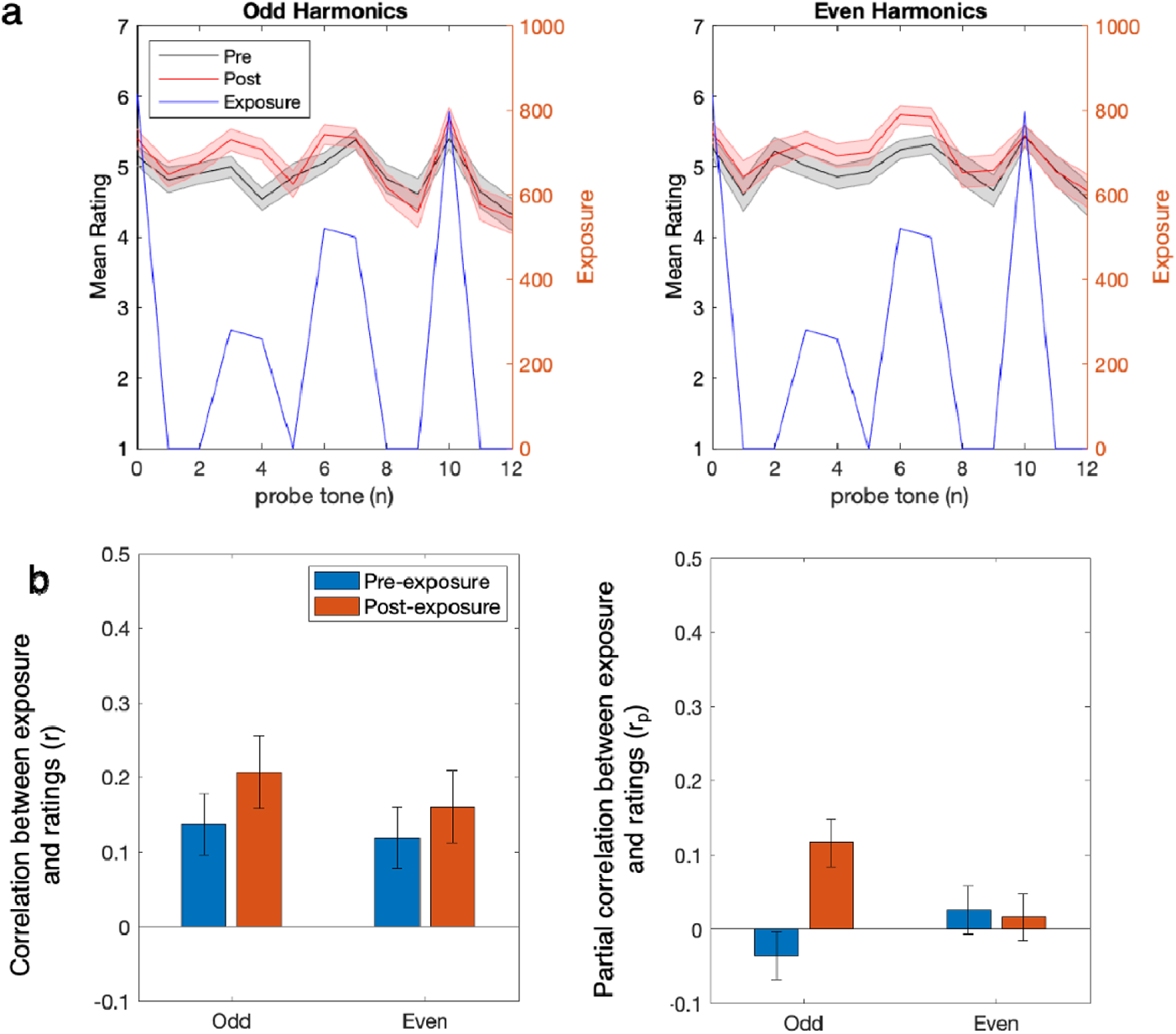
Results from odd and even harmonics statistical learning study. **a)** Probe tone ratings for participants from the odd harmonics condition (left) and the even harmonics condition (right). X-axis represents the probe tone in steps along the BP scale; primary Y-axis shows mean rating on the scale of 1 to 7; secondary Y-axis shows frequency of exposure. Pre-exposure ratings are in black and post-exposure ratings are in red. The exposure profile is shown in blue, and its axis is on the right of each plot. Shaded error bars represent ±1 between-subject standard error. **b)** Correlations (left) and partial correlations (right) between probe-tone ratings and exposure frequencies. Error bars show between-subject standard error.

To check whether the Pearson correlations and partial correlations were normally distributed, a one-sample Kolmogorov-Smirnov test was run on the pre-exposure correlations, the post-exposure correlations, the pre-exposure partial correlations, and the post-exposure partial correlations. All K-S tests were not significant (all p’s > .2), confirming that the correlations did not deviate from normal distribution. These correlations and partial correlations were then entered as dependent variables into a mixed effects multivariate ANOVA, with the within-subjects factor of exposure (pre- vs. post-exposure) and the between-subjects factor of condition (odd vs. even harmonics). Results of this model are shown in Table 3. Results showed a significant within-subjects effect of exposure (F(2,124) = 3.52, p = .033, partial η^2^ = .054), confirming a difference between pre- and post-exposure ratings. The effect of condition was marginally significant (F(2,124) = 2.80, p = 0.064, partial η^2^ = .043). Importantly, there was a significant exposure by condition interaction (F(2,124) = 6.90, p = .001, partial η^2^ = 0.1, confirming the results visualized in Figure 2b that odd harmonics better aided statistical sensitivity and learning of the BP scale.

**Table 3.** Results from Experiment 2. **a)** Mixed effects MANOVA. **b)** Two-tailed t-tests of z-transformed pre- and post-exposure partial correlations against the chance level of zero. * denotes significant deviation at p < .05 from chance level of 0, two-tailed t-test.

| a) Effect | | | F | Hypothesis df | Error df | p | Partial $\eta^2$ | |
| --- | --- | --- | --- | --- | --- | --- | --- | --- |
| Between Subjects | Intercept |  | 30.97 | 2 | 124 | <.001 | 0.333 |  |
|  | Condition |  | 2.80 | 2 | 124 | 0.064 | 0.043 |  |
| Within Subjects | Exposure |  | 3.52 | 2 | 124 | 0.033 | 0.054 |  |
|  | Exposure * Condition |  | 6.90 | 2 | 124 | 0.001 | 0.1 |  |
| b) One-sample t-test against test value = 0 |  |  |  |  |  | 95% Confidence Interval of the Difference |  |  |
|  | Exposure | t | df | p (2-sided) | Mean Difference | Lower | Upper | Cohen's d |
| Even | Pre | 0.8 | 61 | 0.43 | 0.03 | -0.04 | 0.09 | 0.10 |
|  | Post | 0.45 | 61 | 0.66 | 0.02 | -0.06 | 0.09 | 0.06 |
| Odd | Pre | -1.11 | 64 | 0.27 | -0.04 | -0.10 | 0.03 | -0.14 |
|  | Post | 3.61 | 64 | <.001* | 0.12 | 0.05 | 0.18 | 0.45 |

Having observed a significant effect of exposure and a significant exposure by condition interaction, because part of this experiment was done online, we further asked whether the testing context (in-person vs. online) affected learning. We ran another repeated-measures multivariate ANOVA where testing context was incorporated as a covariate, in addition to the between-subjects factor of condition. Results showed a significant interaction of testing context with exposure (F(2,123) = 7.26, p = .001, partial η^2^ = 0.11). To explore this, a follow-up analysis separately tested the effect of exposure and the interaction of exposure-by-condition for online and in-person cohorts. Table 4 shows univariate tests of the main effect of exposure and the exposure-by-condition interaction for correlations and partial correlations, separately for online and in-person testing contexts, and Figure 6 compares correlations and partial correlations between in-person and online testing contexts. Results show that in-person testing yielded higher correlations and partial correlations (Figure 6). Participants from both online and in-person testing contexts showed significant effects of exposure. Online participants showed a significant exposure by condition interaction, but this was not significant in the much smaller sample of in-person participants. The effect size (partial η^2^) was higher for in-person testing for the effect of exposure, but was comparable between in-person and online for the exposure-by-condition interaction (Table 4).

**Figure 6.**
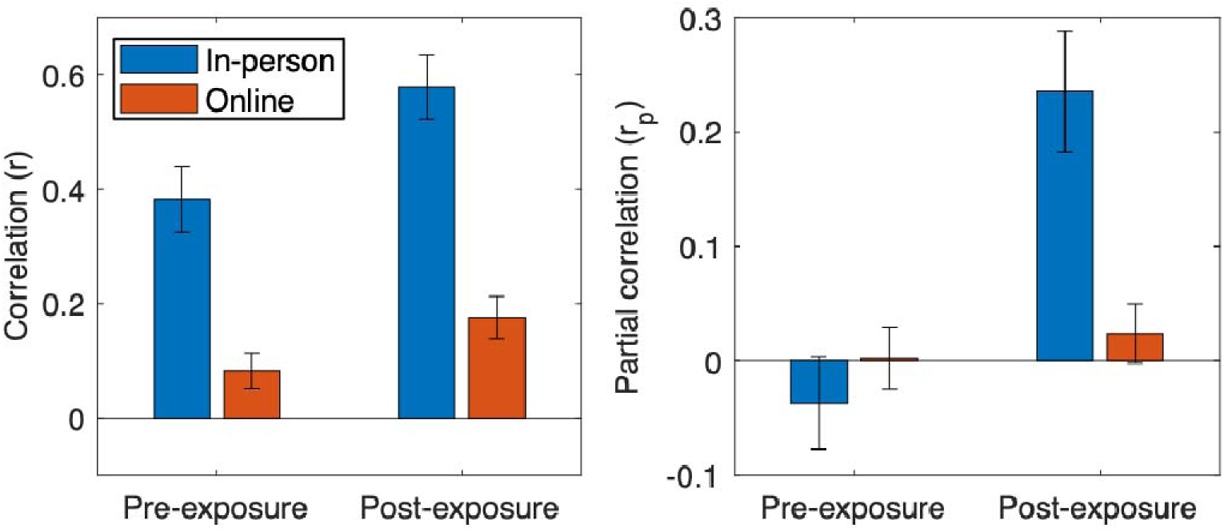
Partial correlations comparing in-person and online testing contexts from Experiment 2.

**Table 4.**
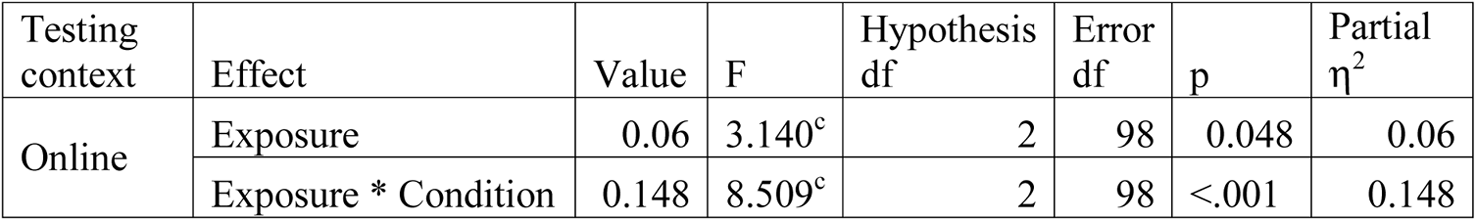
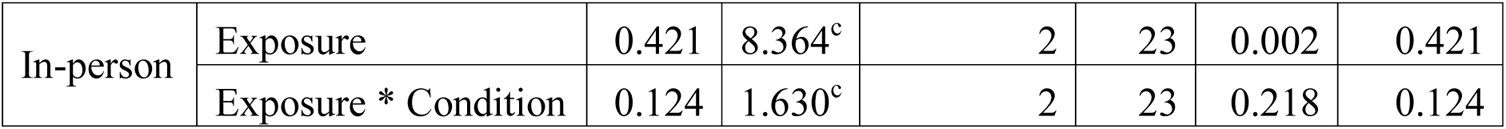
Main effects and interactions from Experiment 2 separately for online and in-person testing cohorts.

### 4.3 Discussion

Results from Experiment 2 showed that participants were better able to learn the statistical structure of the BP scale following exposure in odd harmonics, compared to exposure in even harmonics. The contribution of spectral information to statistical learning is more clearly shown using the larger sample size and the simpler two-group design of Experiment 2, which showed a group-by-exposure interaction to confirm better learning in the odd harmonics group. While there was a significant effect of exposure overall, partial correlations showed that only the odd harmonics group was able to learn. The even harmonics group did not learn significantly from exposure when the effects of the melody used to obtain the ratings were partialled out.

Some part of the lower effect size of learning overall may be explained by the fact that more than half of our participants completed the experiment online on their own. We tried to mitigate the possible effects of inattention during online studies by confirming that participants were using headphones using a headphone screening routine, and providing the task instructions during the exposure phase which specified that participants should not remove their headphones and should not talk or listen to other sounds during the half-hour exposure. Inspecting the work time that it took to complete the online experiment showed that all participants took more than 35 minutes to complete the experiment, suggesting that participants did indeed remain online for at least the duration of the exposure. Still, it was more difficult to ensure that participants indeed listened attentively during the exposure in this less controlled environment, which may explain the lower mean correlations in Experiment 2 than Experiment 1. Nevertheless, the online testing environment could not explain the predicted and observed interaction between group and exposure, as both groups were tested in the same environments and yet the odd harmonics group learned better. This superior learning in the odd harmonics group could only be explained by the congruency between spectral information and scale structure. Thus the current experiment provides strong evidence in support of the influence of spectral information on learning of a musical scale.

## 5 General Discussion

Musical scales around the world are built around whole-integer mathematical ratios in frequency, which are perceived as consonant, but there are cultural and training-related differences in preference for consonance (McDermott, Lehr, & Oxenham, 2010; McDermott, Schultz, Undurraga, & Godoy, 2016), and we do not know why consonant intervals are relatively important in musical scales across many cultures. Some theories posit that musical scale structures reflect statistical properties of periodic sounds in the environment, such as speech sounds (Bowling & Purves, 2015). Support for this comes from the association between speech sounds and consonant pitch intervals, but until now this has been mostly correlational evidence. Here, using a new musical system to which participants are exposed for the first time, we test the causal relationship between sound spectrum and learning of musical structure. Experiment 1 showed that all participants who received exposure were able to learn the BP scale, but tritave complex tones with partials congruent with the scale structure appeared to lead to better learning, although the exposure by condition interaction was not statistically significant. This was extended in Experiment 2, in which stimuli contained more spectral information overall, as the stimuli in Experiment 2 contained complex tones of seven frequency components in addition to the fundamental, compared to Experiment 1 which included only four frequency components in addition to the spectral centroid. Experiment 2 showed that odd harmonics led to better learning than even harmonics, providing more support for the role of spectral information on statistical learning.

Participants were better at learning the BP scale when they heard melodies in the BP scale presented in a timbre that was consistent with BP scale structure. Results show a relationship between timbre –– specifically the spacing between adjacent harmonics –– and the learning of scale structure, thus providing the first support for the role of sound spectrum in the statistical learning of music.

Learning of scale structure was quantified using the exposure-related change in correlation between subjective ratings from the probe-tone method and the distribution of event probabilities of exposure. Probe-tone methodology has shown sensitivity to musical scale structure (Krumhansl, 1990). Here, by comparing probe-tone profiles before and after exposure to tone sequences in a new musical system, we can capture new musical knowledge as it emerges for the first time as a result of ongoing statistical learning from short-term exposure. Results converge with existing literature on statistical learning to demonstrate that the human brain is a flexible learner that adapts rapidly to form predictions for the frequencies and probabilities of sounds in the environment (Daikoku, Yatomi, & Yumoto, 2017; Jonaitis & Saffran, 2009; Pearce et al., 2009; Saffran et al., 1999).

While other studies have conceptualized statistical learning as the sensitivity to transitional probabilities (e.g. Saffran et al, 1996), which are first-order probabilities of an event given its previous event, here we conceptualize the sensitivity to scale structure as zero-order probability, or the distribution of frequency of occurrence across different pitches along the musical scale. This sensitivity to scale structure underlies musical tonality and is best captured behaviorally using the probe-tone method. Previous evidence from electrophysiological recordings (Loui et al., 2009) also support the idea that musical harmonies in the BP scale can be rapidly acquired via statistical learning, thus converging with the present results.

In both experiments, pre-exposure probe-tone ratings showed significant correlation with the scale structure, however these dropped to chance levels after partialling out the effect of the tone sequence used to obtain the probe-tone ratings. This suggests that contextual information played a role in these ratings. Since the Tritave complex tones and the odd harmonics are congruent whereas Octave complex tones and even harmonics are incongruent with the tritave-based tuning system, more accurate ratings for congruent timbres than for incongruent timbres suggest that the timbre, specifically the spectral content of sounds, affected participants’ sensitivity to frequencies of tones in their input. While the study of timbre is often manipulated by varying musical instruments (Marozeau, de Cheveigne, McAdams, & Winsberg, 2003; Menon et al., 2002; Shahin, Roberts, Chau, Trainor, & Miller, 2008), here spectral content was varied systematically by changing the arrangement of frequency components in pitches, resulting in complex tones that vary in the spread of energy across the frequency spectrum while keeping other acoustic features constant. This more controlled method ensures that spectral content can be manipulated independently of temporal content and spectrotemporal flux, which are two of the other features that contribute to the overall percept of timbre (McAdams, 2013). Given that Experiments 1 and 2 differ in sample size as well as in spectral manipulations, it remains unclear whether the null effects in Experiment 1 reflect power issues related to the sample size, or to a genuine lack of effect with this particular manipulation. While future experiments that test learning with complex tones with a larger sample size will be informative, we also note that Experiment 2 resulted in a larger effect size (<u>partial η^2^ = 0.1, compared to 0.044 in Experiment 1) of the exposure by condition interaction, suggesting that the larger amount of spectral information in Experiment 2, by including multiple harmonics, may contribute to learning outcomes over-and-above methodological considerations of statistical power.</u>

Although the present study taps into a fundamental aspect of musical ability, participants were unselected for musical training, as previous studies on BP scale learning had shown that statistical learning of the new musical system did not interact with musical training (Loui et al., 2010). Indeed, the results show robust learning as indicated by increased correlations over time, as well as sensitivity to context as indicated by a decrease in correlation scores when effects of context were partialled out. Rather than reflecting a music-specific ability, performance on the rating tasks here may reflect more domain-general learning abilities that also underlie the input-based acquisition of other materials such as speech and language, and environmental sounds more generally. Thus, results converge with findings from the domain of speech and language acquisition, where timbre is found to play a role in mother-infant communication (Piazza, Iordan, & Lew-Williams, 2017).

In this broader context, our results contribute to a growing body of evidence on the importance of timbre, specifically spectral information, as a crucial source of input in forming our schemas for speech, music, and the auditory environment more generally. While the current results only tested sensitivity to scale structure by assessing learning of the event frequencies of pitches in the scale, future studies could test the effectiveness of timbre for learning higher-order statistical structure. For example, it would be interesting for future studies to see whether these results could generalize to first-order transitional probabilities or non-adjacent dependencies (Newport & Aslin, 2004), that have been observed in tone sequences in Western pitch categories (Creel, Newport, & Aslin, 2004). Furthermore, the observed relationship between spectral information and musical structure is implicit in one argument for the naturalness of harmony in Western music, which states that the statistical structure of naturally-occurring periodic signals, such as speech, can predict perceived pitch and other musical phenomena such as the relative stability of consonant musical intervals in the chromatic scale (Schwartz, Howe, & Purves, 2003). Support for this comes from findings of similarities between the periodicity of speech sounds and perceived pitch (Schwartz & Purves, 2004) and also in covariations between speech and musical pitch patterns across cultures (Han, Sundararajan, Bowling, Lake, & Purves, 2011). While these findings provide correlative evidence for a relationship between environmental sounds and musical structures, a causal test of this relationship is challenging with naturally-occurring speech sounds and musical structures, because of the aforementioned inherent difficulty in teasing apart prior long-term knowledge from knowledge acquired from the current auditory environment. The present results fill that gap by offering a new musical system as a window into this relationship between sound spectrum and scale structure. Future studies may test whether people who have more experiences with naturally-occurring sounds that have odd harmonics may be better at learning the BP scale, thus offering an even closer test to the hypothesis by Purves et al of the relationship between naturally-occurring periodic signals and musical structure.

There are some limitations in the present work. Firstly, Experiment 1 had a relatively small sample size which was determined from previous studies, but could have been underpowered. While this could have contributed to the only trend-level significance of the effect of condition on ratings, rather than arbitrarily increasing the sample size, we used the results of Experiment 1 to power Experiment 2. In doing so we also strengthened the manipulation by changing the complex tones from logarithmic spacing to odd / even harmonic spacing, thus increasing the spectral content by increasing the number of harmonic components that could fit in the spectrum. Both the increased sample size and the stronger spectral manipulation could have contributed to the exposure-by-condition interaction in Experiment 2. An additional complication was that due to the Covid-19 pandemic, the majority of Experiment 2 was moved online, making it more difficult to enforce that participants were listening during the exposure period. Follow-up analyses showed that in-person participants performed better than online participants as shown by higher correlations and partial correlations. Both in-person and online participants were able to learn from exposure. Although only online participants showed a statistically significant exposure-by-condition interaction, effect sizes of this interaction were comparable between in-person and online participants, suggesting that the lack of a significant interaction in the in-person participants may be explained by the much smaller sample size of the in-person cohort. Taken together, while there are limitations that arise from sample size and testing context, these challenges could not explain the overall finding that harmonics that were congruent with the scale aided the learning of scale structure.

Our interpretation of these findings borrows from the free energy principle, the notion that the central nervous system aims to minimize prediction error by dynamically sampling the environment to continuously adapt its prior expectations in a context-sensitive fashion (Friston, Kilner, & Harrison, 2006). In music cognition specifically, learning music entails reducing uncertainty by forming and testing predictions continuously based on context (Friston & Friston, 2013; Hansen & Pearce, 2014; Koelsch, Vuust, & Friston, 2018). The BP scale system offers a new context in which the cognitive system must learn to form novel expectations in order to reduce uncertainty. The information that the cognitive system uses must come from perceptual input, which in the auditory domain generally starts with the breakdown of complex stimuli into auditory filters (Hafter, Sarampalis, & Loui, 2007). Studies on auditory attention have shown that the cognitive system selectively sharpens these auditory filters (Schlauch & Hafter, 1991), even attending to auditory filters in the absence of direct acoustic stimulation (Hafter, Schlauch, & Tang, 1993). Here, when faced with the new context of the BP scale, in order to reduce uncertainty the central nervous system must monitor its perceptual input across all auditory filters, effectively listening for spectral components that share auditory filters that are more likely to be stimulated over the course of exposure. Successful monitoring or attentional sampling of the relevant auditory filters enables better prediction of acoustic events in the novel auditory environment of the BP scale. Future studies, perhaps using computational modeling and electrophysiological tools, may explore the neural mechanisms of how the selective sampling of auditory filters across the frequency spectrum may accumulate over the course of exposure, thus using the spectral distribution of timbre as a cue for statistical learning to reduce uncertainty in the perceptual environment.

In sum, the present work uses a new music system to shed light on psychological and biological theories of learning. By reimagining the conventions of musical systems, we can start to answer novel questions about the extent to which our minds derive structure from exposure to sounds within our environment.

## 6 Acknowledgements

This work was supported by the National Science Foundation (CAREER award # 1945436) and the Grammy Foundation. I thank three anonymous reviewers for helpful comments, and Ervin Hafter and the late David Wessel for helpful comments during the early-stage conceptualization of this work.

